# Comprehensive Network Analysis of miRNA Biogenesis Proteins and Their Ligand Interaction Sites to Identify Novel Targets for Cancer Therapeutics

**DOI:** 10.1101/2024.12.04.626869

**Authors:** Hazem Almhanna, Hassan Hachim Naser, Arun HS Kumar

## Abstract

A network analysis of canonical microRNA (miRNA) biogenesis identified DROSHA, Exportin-5, and DICER1 as essential proteins for both precursor and mature miRNA processing. The analysis revealed strong interactions between these proteins and others involved in miRNA biogenesis, suggesting a complex regulatory network. Ligand binding sites on these key proteins were identified, suggesting potential targets for therapeutic intervention. Our findings indicate that modulating miRNA biogenesis through these proteins could influence cellular protein production and function, providing a promising avenue for developing advanced therapy medicinal products (ATMPs) to impact protein expression in diseases such as cancer.

## Introduction

Recent studies confirmed that mature microRNAs (miRNAs) are small non-coding RNA molecules (typically about 21-25 nucleotides in length) expressed in animal, plants cells, unicellular organisms, and some viruses and play a crucial role in regulating gene expression.(1) The miRNAs are a type of non-coding RNA, and do not associate with proteins synthesis.(2) However, miRNAs control gene activity by binding to specific messenger RNA (mRNA) molecules. These mRNA molecules carry the genetic information necessary for protein synthesis.

During translation, the process of converting mRNA into protein, miRNAs come into action.(3) Mature miRNAs are produced in a multi-step process. First, a longer precursor molecule, called pri-miRNA, is transcribed from DNA. This pri-miRNA is then processed by enzymes into a shorter hairpin-shaped structure called pre-miRNA. Finally, the pre-miRNA is further processed to generate the mature miRNA.(4, 5) The mature miRNA formed can selectively bind to specific mRNA molecules and typically bind to the 3′ untranslated region (UTR) of mRNAs.(6, 7) When a miRNA binds to an mRNA, it prevents the mRNA from being translated into protein or causes its degradation, consequently, miRNAs can fine-tune gene expression by controlling the amount of protein produced from certain genes.(8-10) Numerous studies have confirmed that when a miRNA binds to an mRNA molecule, it can either block the mRNA from being translated into a protein or accelerate its breakdown, leading to a reduction in protein expression.(11, 12) miRNAs are involved in various biological processes, including development, cell differentiation, apoptosis, and immune response. They also regulate gene expression by targeting specific mRNAs that have complementary sequences to the miRNA, usually within the 3’ untranslated region (UTR) of the mRNA.(13) Accordingly, the binding of miRNAs to their target mRNAs is mediated by the RNA-induced silencing complex (RISC), which contains proteins that help stabilize the miRNA-mRNA interaction and facilitate mRNA degradation or translational repression.(14, 15) Importantly, miRNAs have been implicated in numerous diseases, including cancer, cardiovascular diseases, neurodegenerative disorders, and immune-related disorders.(16-18) Therefore, dysregulation of specific miRNAs can contribute to disease progression by altering the expression of genes involved in disease pathways.(19) Recent studies confirmed that stability and presence of the miRNAs in various body fluids suggested it as potential biomarkers for diagnostic, prognostic, and therapeutic purposes.(1, 20-22) There are different types of miRNAs based on their origin and function. Accordingly, canonical miRNAs are the most common type of miRNAs that transcribed from DNA in the cell’s nucleus and then processed into mature miRNAs, and regulate gene expression by binding to mRNA molecules and preventing them from being translated into proteins.(23) This hairpin pre-miRNA is generated from introns and cleaved by DROSHA, a nuclease of the RNase III family. The pre-miRNA is then exported into the cytoplasm via EXPORTIN-5. Subsequently, it undergoes processing by DICER, another RNAse III endoribonuclease, before becoming a mature miRNA.(24)

Also, another special type of miRNA is non-canonical hairpin miRNAs called Mirtrons which are derived from introns through splicing and subsequent debranching.(25) Introns are non-coding regions within genes. The biogenesis of miRNAs and mirtrons differs from that of canonical miRNAs. Mirtrons are spliced out of the mRNA molecule without passing through the DROSHA and DICER processing steps. They are then processed into mature miRNAs, which can regulate gene expression. Mirtrons possess a 3′ terminal guanyl nucleotide, which stimulates Tailor-mediated uridylation and subsequent decay.(26) Hairpin pre-miRNAs undergo cleavage during the Drosha cleavage step of the canonical miRNA pathway. Unlike typical canonical miRNAs, they lack a lower stem and basal single-stranded segments.(25, 27) Consequently, miRNA molecules regulate gene expression by influencing their translation into proteins. This plays a significant role in various biological processes, contributing to the complexity and functionality of living organisms. Therefore, this study focused on network analysis and the biological process synthesis of miRNAs to better understand their functions and explore their potential applications in cancer diseases and medicine.

## Material and Methods

The protein sequences of DROSHA (UniProt ID Q9NRR4), Exportin-5 (UniProt ID Q9HAV4), and DICER1 (UniProt ID Q9UPY3) were retrieved in FASTA format from the UniProt database (https://www.uniprot.org/). Three-dimensional (3D) structures of the proteins were downloaded as PDB files. Network analysis of these proteins was performed using the STRING database (https://string-db.org/) and top ten networks were identified. Briefly the protein name was inputted into the search box and homo sapiens was selected from the dropdown menu for “Organisms” and the search button was clicked. From the search results the protein of interest was selected and the “continue” option was clicked to get the top networks of the protein (28, 29).

To assess the number of hydrogen bonds formed between the protein and its networks, the PDB files of the protein pairs of interest were imported to the Chimera software and the number of intermolecular hydrogen bonds formed were observed at 10 Armstrong bond distance. The number of hydrogen bonds show in the output file was noted in the Excel sheet and the 2D image showing the protein pairs with hydrogen bonds was saved as a jpeg file(30-32).

To identify the binding sites on DROSHA, Exportin-5, and DICER1, the PrankWeb: Ligand Binding Site Prediction tool (https://prankweb.cz/) was used. Briefly, the PDB file of the protein was uploaded to the database and submit button was clicked. From the output data the jpeg file showing the binding sites on the protein was saved and the excel sheet showing the details of the binding sites was downloaded for further analysis. The amino acid sequence of the top binding site was figured by matching the amino acid number with the protein sequence in the FASTA format. The amino acid sequence of the binding site was used to generate antisense 3D structure using the Chimera software and the resulting 3D structure was used to assess its interaction with the primary protein by estimating the number of intermolecular hydrogen bonds formed as above(32).

## Results

This study employed network analysis to investigate the protein components and ligand binding sites involved in canonical miRNA biogenesis. Utilizing the UniProt database, three key human proteins associated with both premature and mature miRNA processing were identified. These proteins include: 1) RNase III (DROSHA), responsible for the initial cleavage of miRNA precursors; 2) Exportin-5, facilitating the nuclear export of pre-miRNAs to the cytoplasm; and 3) endoribonuclease Dicer (DICER), involved in the final processing of pre-miRNAs into mature miRNAs.

The network analysis of the ribonuclease III, exportin-5, and endoribonuclease was performed using STRING database. These proteins showed high interactions with 10 top different proteins listed in table 1, 2, and 3, and Figure1. The alpha fold (AF) structure of these proteins were imported onto the Chimera software and the number of inter molecular hydrogen bonds (H-bond) between them at 10 Armstrong (10A) distance was recorded. The binding sites of ribonuclease III, exportin-5 and endoribonuclease DICER were identified utilizing the PrankWeb tool.

**Table1:** This table displayed network analysis for ribonuclease III (DROSHA).

| No. | Predicted Functional Partners | Gene | Score | HB |
| --- | --- | --- | --- | --- |
| 1 | Probable ATP-dependent RNA helicase DDX5 | DDX5 | 0.999 | 8958 |
| 2 | Microprocessor complex subunit DGCR8 | DGCR8 | 0.999 | 7164 |
| 3 | Probable ATP-dependent RNA helicase DDX17 | DDX17 | 0.998 | 10267 |
| 4 | Cellular tumor antigen p53 | TP53 | 0.996 | 6306 |
| 5 | Protein argonaute-2 | AGO2 | 0.995 | 13589 |
| 6 | TAR DNA-binding protein 43 | TARDBP | 0.992 | 6086 |
| 7 | RNA-binding protein FUS | FUS | 0.992 | 5332 |
| 8 | Endoribonuclease Dicer | DICER1 | 0.992 | 18093 |
| 9 | RISC-loading complex subunit TARBP2 | TARBP2 | 0.989 | 6526 |
| 10 | Protein argonaute-1 | AGO1 | 0.989 | 13045 |

HB: number of hydrogen bonds, Score refers to probability of interaction.

**Table2:**
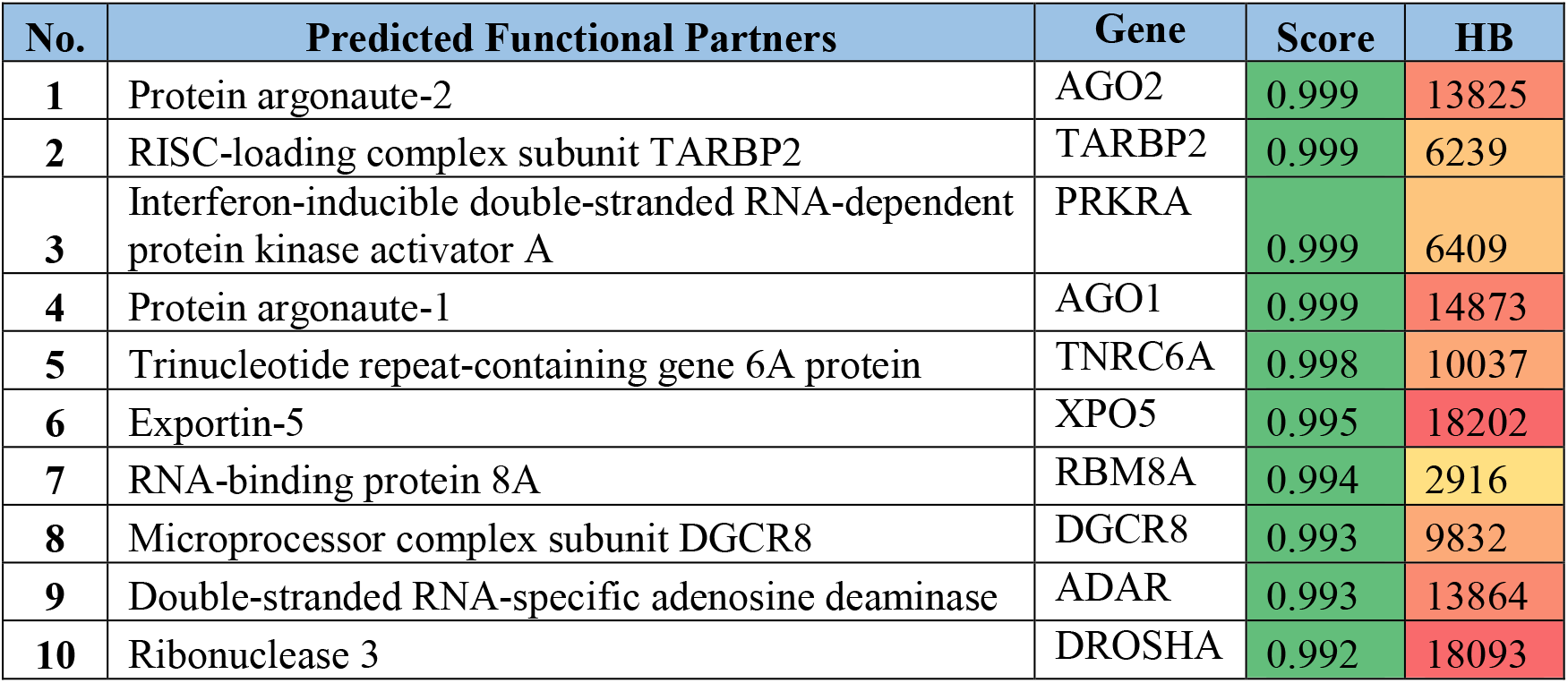
This table displayed network analysis for endoribonuclease DICER (DICER1).

**Table3:**
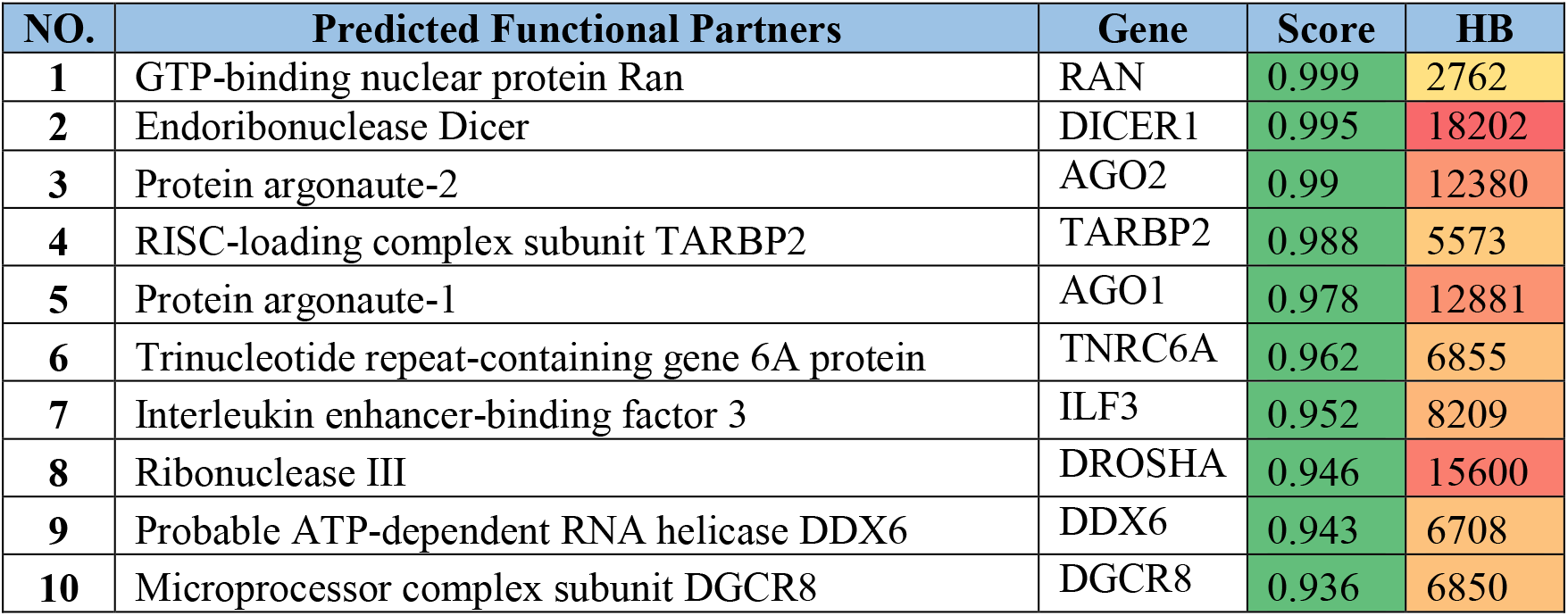
This table displayed network analysis for exportin-5 (EXPORTIN—5).

| **NO.** | **Predicted Functional Partners** | **Gene** | **Score** | **HB** |
| --- | --- | --- | --- | --- |
| **1** | GTP-binding nuclear protein Ran | RAN | 0.999 | 2762 |
| **2** | Endoribonuclease Dicer | DICER1 | 0.995 | 18202 |
| **3** | Protein argonaute-2 | AGO2 | 0.99 | 12380 |
| **4** | RISC-loading complex subunit TARBP2 | TARBP2 | 0.988 | 5573 |
| **5** | Protein argonaute-1 | AGO1 | 0.978 | 12881 |
| **6** | Trinucleotide repeat-containing gene 6A protein | TNRC6A | 0.962 | 6855 |
| **7** | Interleukin enhancer-binding factor 3 | ILF3 | 0.952 | 8209 |
| **8** | Ribonuclease III | DROSHA | 0.946 | 15600 |
| **9** | Probable ATP-dependent RNA helicase DDX6 | DDX6 | 0.943 | 6708 |
| **10** | Microprocessor complex subunit DGCR8 | DGCR8 | 0.936 | 6850 |

The top ten proteins that interacted with DROSHA, DICER1 and Exportin-5 were assessed for inter molecular hydrogen bonds formation. DROSHA displayed high interaction and most hydrogen bonds with DICER1 (18093), AGO2(13589), and AGO1 (13045). While DICER1 showed high interaction and more number of hydrogen bonds with Exportin-5 (18202) DROSHA (18093), AGO1 (14873), AGO2(13825), and ADAR (13864). Exportin-5 showed most number of hydrogen bonds with DICER1 (18202) DROSHA (18093), AGO1 (12881), and AGO2(12380), Shown in table (1,2, 3).

Additionally, the ligand binding sites of ribonucleases III and DICER, as well as exportin-5, were identified as potential targets for development of advance therapy medicinal products (ATMP).

These ATMPs have the potential to interfere with the normal functions of specific proteins, thereby blocking protein production (translation) and subsequent modifications (post-translation). We identified binding sites on three key proteins: ribonuclease III (IRFIPVPRQNFRLSIPY, 341 hydrogen bonds), exportin-5 (FETLSCTN, 61 hydrogen bonds), and endoribonuclease DICER (LYHCTSRMVVSNWLPHTQKAISY, 435 hydrogen bonds) (Figure 1).

**Figure (1):**
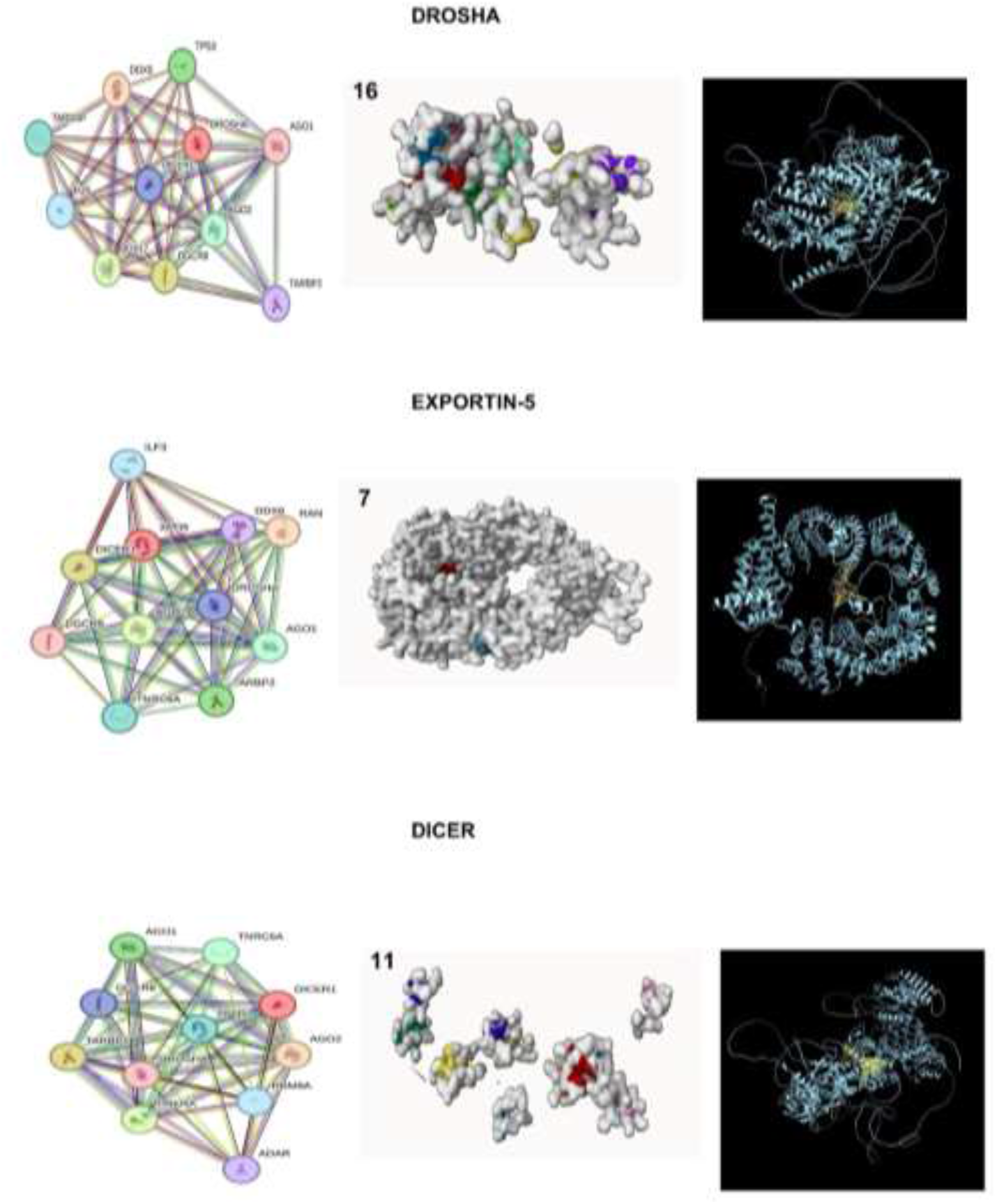
This figure presents a network analysis of DROSHA, Exportin-5, and DICER1, including their ligand binding sites and corresponding antibodies. The coloured values represent the number of identified binding sites for each protein.

**Figure (2):**
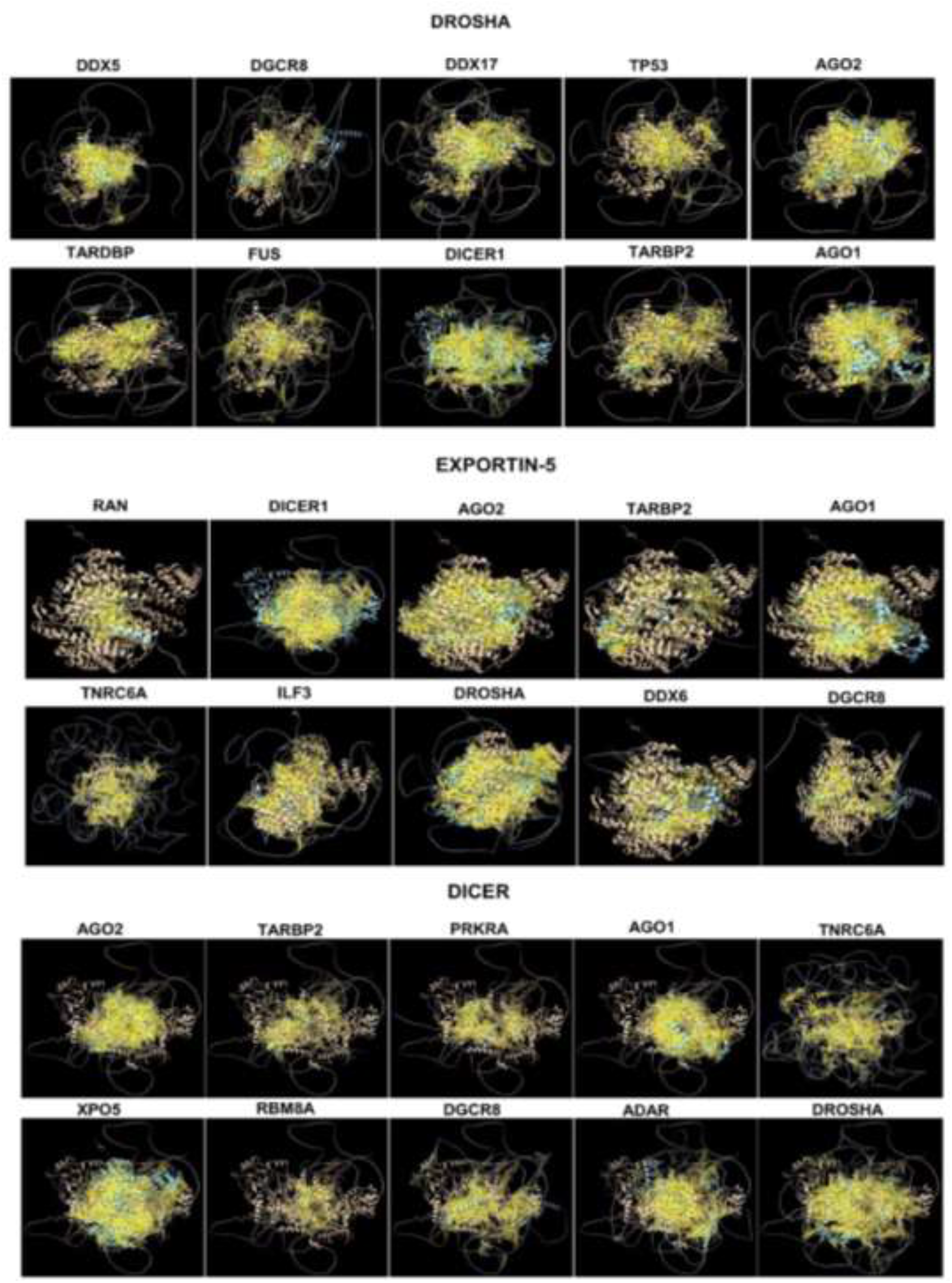
This figure displays a protein-protein interaction of DROSHA, Exportin-5, and DICER1 with their respective network proteins. Yellow lines indicate the number of intermolecular hydrogen bonds.

## Discussion

Recent studies propose that miRNAs do not directly contribute to protein synthesis. However, they can influence the process of coding gene expression of the proteins, leading to either decreased or increased protein production.(12, 33) These research findings suggest that controlling the biogenesis and synthesis processes of miRNAs within cells may affect the activity of cellular proteins and the performance of biological cell functions. The present study utilized network analysis to explore the protein components and ligand binding sites involved in canonical miRNA biogenesis. Our findings highlight the intricate interactions between key proteins, DROSHA, Exportin-5, and DICER, in regulating miRNA production.

Network analysis revealed significant interactions between these proteins and other components within the miRNA biogenesis pathway. The high number of intermolecular hydrogen bonds observed between DROSHA, DICER, and AGO proteins suggest strong physical associations and potential functional cooperativity. These interactions are likely crucial for the efficient processing and transport of miRNAs. Contemporary investigations have found that ribonuclease III (DROSHA), exportin-5, and endoribonuclease DICER are involved in the biogenesis of canonical miRNAs(34-36) and other types of miRNAs. As the biogenesis of miRNAs would likely influence the expression of various types of proteins, any approach to modulate these three regulators of miRNA can have a therapeutic merit. Our study found that DROSHA interacts most strongly with DICER1, followed by AGO2, AGO1, and DDX17 in decreasing order of interaction strength. Hence, DICER1 consistent with the literature can be considered a key enzyme in gene regulation.(37) Previous research has shown that DICER1 plays a vital role in processing precursor RNAs into small interfering RNAs (siRNAs) and microRNAs (miRNAs).(38, 39) These small RNA molecules control gene expression by regulating protein production and are also involved in immune responses and antiviral defence.(40, 41) Overall, these results strongly support the notion that controlling the expression of this enzyme can influence the production of various miRNA molecules. Additionally, these miRNAs are powerful regulators of gene expression and can impact the production of negative proteins, which contribute to the development of malignant cancers or tumors in different organs and tissues.

Our results also demonstrated a robust interaction between DICER1 and Exportin-5. While DICER1 and Exportin-5 strongly interact with each other, our findings suggest a more complex relationship involving DROSHA. Although Exportin-5 was not within the top ten networks of DROSHA or DICER1, both these proteins were within the top ten networks of Exportin-5, with significant number of hydrogen bonds formed between them. Prior research suggests that Exportin-5 binds to pre-miRNAs within the nucleus. This complex then interacts with RanGTP, a small GTPase, to facilitate the export of pre-miRNAs through the nuclear pore complex into the cytoplasm.(42-44) This process influences the expression of one or more specific proteins. Our observations add a new dimension to significant protein-protein interactions between DROSHA, Exportin-5 and DICER1 with potentially involving AGO1 and AGO2 proteins. Both AGO1 and AGO2 were among the top ten networks of DROSHA, DICER1 and Exportin-5. These results support the hypothesis that DROSHA, Exportin-5, and DICER1 play a critical collective role in the assembly of the RNA-induced silencing complex (RISC) during pre- and mature miRNA biogenesis.(45-48). Our analysis of the interplay between DROSHA, DICER1, and Exportin-5 during small interfering RNA (siRNA) and microRNA (miRNA) biogenesis revealed a significant synergy between these proteins. Our findings corroborate the results of other recent studies in this field.(46, 49)

Several studies have reported that miRNAs can act as both cancer promoters and suppressors, and changes in miRNA levels can be used to develop new cancer treatments by targeting mRNAs or their regulators(50-52). In this context, the identification of ligand binding sites on ribonuclease III, exportin-5, and endoribonuclease DICER in this study, offers promising avenues for the development of advanced therapy medicinal products (ATMPs). Targeting these sites with specific ligands could potentially disrupt the normal functions of these proteins, leading to altered miRNA production and subsequent impacts on protein expression. Our results suggest that controlling miRNA biogenesis through these proteins may have significant implications for various biological processes. By interfering with miRNA production, it may be possible to modulate the expression of downstream target genes, thereby influencing cellular function and potentially impacting disease states. However, further research is needed to fully elucidate the precise mechanisms by which these proteins interact and regulate miRNA biogenesis. Additionally, the development of ATMPs targeting these ligand binding sites requires careful consideration of potential off-target effects and selectivity.

In summary, our findings advance our comprehension of miRNA biogenesis and its role in cellular function. By clarifying the interconnectedness of DROSHA, DICER1, and Exportin-5, we provide the groundwork for further exploration of miRNA regulatory mechanisms. Finally, this data may cover the way for innovative therapies targeting miRNAs to treat diseases like cancer by modulating gene expression.

In conclusion, this study provides valuable insights into the protein network underlying canonical miRNA biogenesis. The identification of key proteins and ligand binding sites offers promising opportunities for therapeutic intervention. Future research should focus on further characterizing these interactions and exploring their potential applications in disease treatment.

## Interest of Conflict

**None**

## Notes

### Competing Interest Statement

The authors have declared no competing interest.

